# Soluble LIGHT (TNFSF14) activates endothelial cells, thereby priming the first vessel-occlusive events in acute sickle cell disease

**DOI:** 10.1101/2023.06.08.541088

**Authors:** Philippe Chadebech, Kim-Anh Nguyen-Peyre, Gaétana Di Liberto, Gwellaouen Bodivit, France Pirenne, Pablo Bartolucci

## Abstract

In sickle cell disease (SCD), the red blood cells carry a mutated form of hemoglobin (HbS) leading to altered shape and deformability. The mutation causes abnormal hemorheological properties, mechanical hemolysis, and adhesion. The chronic vascular inflammation observed in SCD and hemolysis-related endothelium activation may trigger the vaso-occlusion of blood vessels.

The prothrombotic and pro-inflammatory LIGHT/TNFSF14 is a tumor necrosis factor (TNF)- superfamily cytokine implicated in various inflammatory diseases. It is expressed by various immune cells and is considered an actor in T cell-mediated immunity and immune cell recruitment. LIGHT has also been shown to activate endothelial cells (ECs) strongly. LIGHT levels are high in the plasma of SCD patients, and platelets are a major source of its circulating form.

We studied a cohort of 82 homozygous adult patients with SCD (*n*=108 samples) to determine whether LIGHT levels were linked to the clinical state of patients included in the ‘*Basal*’ steady state or during an ‘*Acute*’ crisis. Soluble LIGHT levels were high in the plasma of SCD patients during acute phases of the disease, particularly during painful occlusive crises. LIGHT levels were associated with Hb levels and inflammatory markers (mainly interferon-γ and tumor necrosis factor-α, specifically in acute SCD patients). Our findings confirm that LIGHT is a strong activator of cultured ECs, inducing a type II inflammatory cytokine profile and the expression of adhesion molecules. Using a physiological flow adhesion test on biochips, we showed that the LIGHT-induced activation of ECs led to the adhesion of both sickle platelets (but not their AA counterparts), and in a less extend sickle RBCs to activated HUVECs, potentially constituting the first step in vaso-occlusion. Indeed, the pretreatment of HUVECs with neutralizing polyclonal Abs against LIGHT, but not the non-specific counterpart, showed a reversal of both the inflammation process activated by LIGHT treatment and platelet adhesion to endothelial cells.

Soluble LIGHT appears to be a promising therapeutic target for preventing adverse occlusive events in SCD through the blockade of its receptor, to prevent the adhesion of blood cell components to the endothelium. Future studies should consider whether soluble LIGHT contributes to other clinical complications in SCD.

**HIGHLIGHTS:**

- In patients with SCD, plasma LIGHT is mainly secreted during acute phases, including VOCs.
- Both Hb levels and IFNγ levels are correlated with plasma LIGHT with SCD patients in acute phase disease.
- The LIGHT-induced activation of endothelial cells leads to the flow adherence of both sickle (SS) platelets and red blood cells.
- Both endothelial priming by LIGHT and platelet adherence are abolished by anti-LIGHT polyclonal Abs.

## INTRODUCTION

Sickle cell disease (SCD) is a widespread inherited disorder due to a single mutation of the hemoglobin (Hb) gene (HbS)^1^. This mutation alters the characteristics of Hb S-carrying red blood cells (RBCs), modifying their shape and hemorheological properties. These changes cause mechanical hemolysis, but also adherence to blood cell components and the vasculature^2^. HbS-carrying RBCs are responsible for occlusion and organ damage, in association with inflammatory activation of the endothelium or platelets. The transfusion of AA RBC concentrates obtained from donors remains the cornerstone of the standard of care for SCD, and for treating or preventing cerebral vasculopathy and painful events^3^. Vaso-occlusive crisis (VOC) is the principal reason for SCD patient hospitalization^4^; these crises are often complicated by acute chest syndromes (ACS)^5, 6^. Vessel occlusion has been described in SCD patients, with increases in leukocyte and reticulocyte counts, lactate dehydrogenase (LDH) levels and inflammatory markers in patients^7, 8^; such occlusion can lead to tissue infarction and perpetuation of vascular inflammation.

The LIGHT (TNFSF14, CD258, HVEM-L) factor belongs to the tumor necrosis factor (TNF)-superfamily and is expressed by various types of immune cells^9, 10^. LIGHT is associated with platelet membranes and is released during platelet activation^11^. Activated immune cells (including T lymphocytes and natural killer (NK) cells) can also produce large amounts of soluble LIGHT^12–15^. Soluble LIGHT may therefore be considered a potent actor in T cell-mediated immunity. A role for LIGHT in the recruitment of T regulator (T Reg) cells has also been documented^15, 16^. Soluble LIGHT is clinically involved in chronic heart failure^17^ and chronic infections^13, 15^, coronary heart disease and atherogenesis^10, 11, 18^; it may also be involved in wound healing^12^ or susceptibility to *Leishmania* infections^16^. The preclinical development, over the last decade, of methods for delivering LIGHT to tumors has raised hopes of new approaches for enhancing treatment in the field of cancer immunotherapy^19, 20^.

Plasma LIGHT levels are higher in SCD patients than in controls. LIGHT expression levels are also high on the membranes of SCD platelets, confirming that these cells are the major source of circulating LIGHT in this disease^8^. Plasma LIGHT levels in SCD are known to be biologically correlated with the levels of LDH and certain inflammatory markers (CD40-L, IL-8 and CD54/ICAM-1). Furthermore, higher LIGHT concentrations have been implicated in the pathophysiology of SCD through their association with high tricuspid regurgitant velocity (>2.5 m/s) in patients. *In vitro*, both biologically derived LIGHT and human recombinant (hr) LIGHT induce a pro-inflammatory state in vascular endothelial cells (ECs), with an upregulation of adhesion molecules and release of chemokines^11^ through the activation of NF-κB-dependent genes^21^. Circulating LIGHT has a potent activating effect on ECs and may, therefore, contribute to endothelium activation and inflammation in SCD^8^. Finally, the membrane-associated LIGHT on platelets can mediate platelet adherence to the endothelium^10^.

We investigated the associations between various clinical contexts in SCD and soluble LIGHT levels. We assessed LIGHT concentrations in plasma samples from a cohort of selected patients in different clinical phases of the disease. The role of LIGHT in promoting HUVEC activation was also investigated, together with the adherence of RBCs and platelets to ECs, as such mechanism could trigger vaso occlusions in SCD patients. Finally, we assessed the associations between plasma LIGHT concentration and the biological/hematological parameters of SCD patients.

## MATERIALS AND METHODS

Full details of blood products, primary HUVEC isolation and *in vitro* culture on biochips are provided in the Supplementary Methods section.

### SCD patients and controls

We studied a cohort of 82 adult homozygous HbS patients (with a total of *n* = 108 blood samples collected) during transfusion follow-up. Samples were assigned to the following categories: patients with steady-state SCD and no recent pain or transfusion (‘*Basal*’; *n* = 28) and patients with painful acute-phase SCD requiring transfusion (‘*Acute*’; *n*=33). Patients experiencing a post-transfusion delayed hemolytic reaction were identified separately (‘*DHTR*’, *n* = 21) and, when available, their pretransfusion samples were also identified (‘*pre-DHTR*’; *n* = 15). Finally, we selected 21 volunteer blood donors of African descent as a control group.

### RBC collection and preparation

Fresh RBCs were isolated from blood samples by centrifugation at 2880 x *g*, for 10 min at +20 °C. The plasma was decanted off and the RBCs were washed in 0.9 % NaCl and collected as a pellet by centrifugation.

### Primary HUVECs: isolation and culture

Human vascular endothelial cells (HUVECs) were isolated from fresh umbilical cord veins obtained from the gynecology-obstetrics department, as previously described^22^. Primary cells obtained were detached by incubation with 0.5 mg/mL collagenase and cultured in EGM-2 medium. They were used at passages 4 to 6 in all experiments.

### Detection of HUVEC activation and adherence assays

#### # flow cytometry

HUVECs cultured in plastic flasks were left untreated or were treated with hrLIGHT (50, 100, 200 ng/mL; R&D Systems Bio-techne, Minneapolis, MN, USA) diluted in 0.2 % BSA in PBS, for 4, 8 or 20 hours at +37 °C. TNFα (20 ng/mL; R&D Systems) was used as a positive control for all time points. Culture supernatants were frozen at −80 °C for further analysis. HUVECs were washed in HBSS Cold PBA/lidocaine (PBS supplemented with 10 g/L albumin, 5 mg/mL lidocaine chlorhydrate, 10 mM EDTA) was added, with gentle shaking, to detach the cells. Cells were rinsed and stained by incubation 15 min. at +20 °C with the following monoclonal antibodies (mAbs) in 0.2 % BSA in PBS: BB515-conjugated anti-CD54/ICAM-1, PE-conjugated anti-vWF, APC-conjugated anti-CD106/VCAM-1, BV421-conjugated E-selectin/CD62-E and BV510-conjugated anti-CD31/PECAM-1 (BD Biosciences). They were fixed by incubation for 15 min with 1X Cell FIX (BD Biosciences) and then analyzed on a FACS Canto II flow cytometer (BD).

#### # fluorescence microscopy

HUVECs cultured in fibronectin-coated µ-Slide 0.4 Luer (Ibidi, Gräfelfing, Germany) were left untreated or were treated with hrLIGHT/ TNFSF14 (200 ng/mL) for 4 hours at +37 °C. Cells were washed and stained, by incubation for 15 minutes at +20 °C with the following antibodies: BB515-conjugated anti-CD54, APC-conjugated anti-CD106 and BV510-conjugated anti-CD31 antibodies. They were then mounted in DAPI-containing medium. Data were collected on an Axio Imager Zeiss Z1 microscope (Carl Zeiss, Marly le Roi, France) and analyzed with Zen software.

### Luminex detection assay

Plasma samples and culture supernatants were tested in a multiplex cytokine assay using Luminex technology (Affymetrix, Vienna, Austria). The activation profile of HUVECs was determined with a panel of 12 selected cytokines, chemokines, and growth factors. For determination of the cytokine profiles of SCD patients, we determined the following soluble plasma factors, as previously described^23^: IL-6, IL-10, IL-27, IFN-γ, TNF-α, ICAM-1, IP-10, MIP-1β and RANTES.

### In vitro microfluidic model for HUVECs and adherence tests

**<u>Step 1</u>**. **Flow culture of HUVECs**: Cells used to seed a fibronectin-coated IBIDI microfluidic device. They were allowed to adhere to the support for two hours (static conditions) and were then subjected to intermittent flow overnight (shear stress: 1 dyne/cm^2^) with the microfluidic KIMA Pump system (Cellix)^24^. **<u>Step 2</u>. HUVEC conditioning**: HUVECs prepared as described in ‘step 1’were then conditioned by perfusing the microfluidic device at a flow rate of 350 µL/min for 4 hours at +37 °C, under an atmosphere containing 5 % CO_2_; the infusion flow rate was adjusted to the physiological level of 1 dyne/cm^2^ shear stress according to fluid viscosity. Where indicated, EGM-2 medium supplemented with hrLIGHT (500 ng/mL), was infused through the system. **<u>Step 3</u>. Adherence assay**: Whole-blood samples from AA donors or SS patients were collected into heparin and then perfused through the system of conditioned HUVECs obtained as in ‘step 2’ for 10 minutes. A washing step, with culture medium at 1 dyne/cm^2^, was then implemented to remove all circulating blood cells. The cells in the IBIDI microfluidic device were stained by incubation for 15 minutes at +20 °C with the following mAbs: BB510-conjugated anti-CD54, BB515-conjugated anti-CD41a or APC-conjugated anti-CD45. They were then mounted in DAPI-containing medium. Data were collected on a Axio Imager Z1 with Zen software. <u>Alternative procedure</u>: RBCs isolated from fresh blood samples, washed and diluted in EGM-2, were used to perfuse the system, for the assessment of functional adhesion to HUVECs. Adherence was measured under flow conditions with a Mirus Evo Nanopump (Tebu-bio, Le Perray-en-Yvelines, France) compatible with the IBIDI flow chambers used^24–26^. RBC adhesion was quantified by phase-contrast microscopy on an Axio Imager Z1 (Carl Zeiss, Marly le Roi, France) equipped with a black and white Axiocam 702 camera and VenaFlux Assay 2.3 analysis software (Cellix). Attached RBCs were counted with ImageJ by a person blind to the patient’s clinical status and treatments, in five representative areas along the centerline of the biochip for each set of conditions.

### Ethics statement

All the patients were included in a prospective single-center observational study: the SCD-TRANSFU cohort^23, 27^. This research was performed in accordance with the Helsinki Declaration. The medical ethics committee of Henri-Mondor Hospital approved this study (CCP 11-047).

### Statistics

Quantitative data are expressed as the arithmetic mean plus or minus the standard deviation (± SD). Statistical analyses were performed by analysis of variance (ANOVA), with the Kruskal-Wallis test for more than two groups, the Wilcoxon signed-rank test for paired observations and Mann Whitney *U* tests, as appropriate. Data were analyzed with GraphPad Prism 6.01 software (La Jolla, CA, USA). Receiver operating characteristic (ROC) cure analyses were performed to determine the potential diagnostic value of plasma LIGHT concentration in SCD patients. Correlations between biological parameters were investigated by simple linear regression and Pearson’s correlation analyses. Values of *p* < 0.05 were considered significant: * *p* < 0.05, ** *p* < 0.01, *** *p* < 0.001, **** *p* < 0.0001.

## RESULTS

### Soluble LIGHT is secreted into the plasma in SCD patients, particularly during acute phases

We first investigated soluble LIGHT levels by ELISA on plasma samples from our cohort of patients (**Figure 1A and Supplemental Fig. 1A**). We confirmed that plasma LIGHT protein concentrations were significantly higher (p = 0.0002) in samples from SCD patients (mean value: 295.7 pg/mL) than in those from healthy AA donors (mean value: 82.55 pg/mL) (**Supplemental Fig. S1A**). ROC curve analyses were also performed to determine the diagnostic value of plasma LIGHT concentration (AUC: 0.8298 [95 % CI: 0.7371 to 0.9225]; *p* < 0.0001). When SCD patients were split into two groups — basal steady-state patients (‘*Basal’*) or painful acute-phase patients (‘*Acute’*) — plasma soluble LIGHT concentrations were found to be significantly higher (*p* = 0.0004) for acute patients (mean values: 206.1 pg/mL for basal steady-state *versus* 459.9 pg/mL for acute painful SCD samples) (**Supplemental Fig. S1C**).

**Figure 1.**
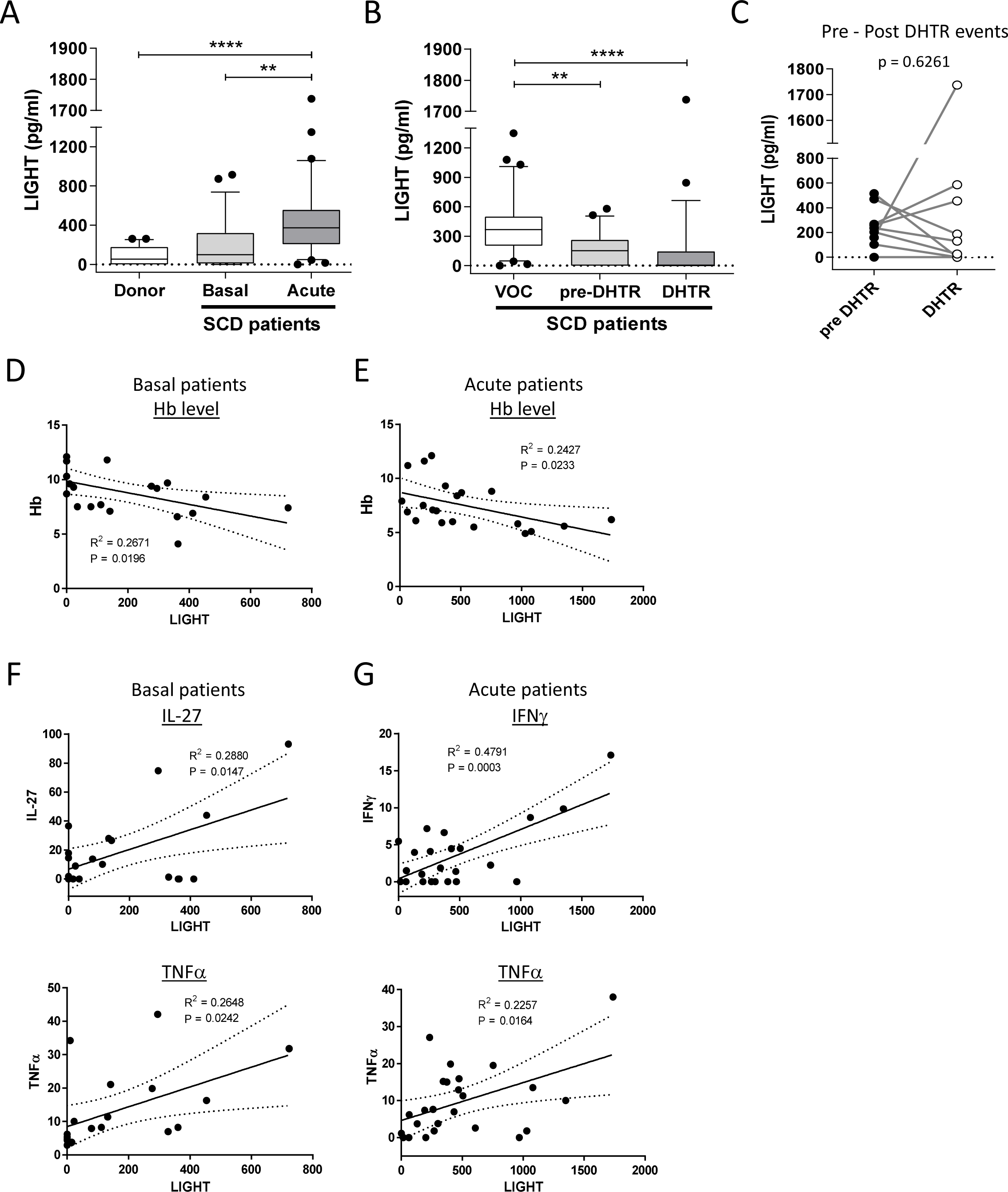
Detection of soluble LIGHT (TNFSF14) factor in plasma samples from SCD patients in different clinical states compared to healthy donors. **(A)** Pre-transfusion plasma samples from SCD patients in a basal steady-state (Basal; *n* = 28; light-gray box) or a painful acute phase (Acute; *n* = 33; gray box) were compared with plasma samples from the control group of blood donors (Donor; *n* = 21; white box). ELISA was performed to quantify soluble LIGHT factor in the various clinical conditions.**(B)** and **(C)** Pre-transfusion plasma samples from patients with painful crisis events leading to a post-transfusion hemolytic reaction (pre-DHTR; *n* = 15; light-gray box) or not leading to such a reaction (VOC; *n* = 32; white box), and from patients sampled at the time of such events (DHTR; *n* = 21; gray box), were subjected to ELISA to quantify the soluble LIGHT present. In **(C)**, LIGHT levels in plasma samples collected before and at the onset of DHTR were compared in pair-wise comparisons. **(D)** and **(E)** show the significant correlation obtained in analyses of the linear regression between Hb levels and LIGHT levels in plasma samples from ‘*Basal*’ and ‘*Acute*’ SCD patients. **(F)** and **(G)** show the significant correlation between LIGHT levels and IL-27 and TNFα concentrations in plasma samples from ‘*Basal*’ SCD patients **(F)** and the significant correlation between LIGHT levels and IFNγ and TNFα levels in plasma samples from ‘*Acute*’ SCD patients **(G)**. Concentrations are shown in box/whisker plots with 10^th^–90^th^ percentile error bars, with dots representing extreme values. ** and **** indicate values of *p*<0.01 and 0.0001, respectively (ANOVA).

Our research work focuses on hemolytic transfusion reactions, particularly those occurring with a time lag after a transfusion event (11delayed hemolytic transfusion reactions11, DHTR) in the context of SCD. We recently published data providing the first evidence of a link between changes in cytokine levels and the occurrence of DHTR^23^. We therefore decided to quantify plasma LIGHT levels and to compare them between the pretransfusion samples from acute-phase SCD patients (the same patients as in **Figure 1A**) classified into three categories: acute-phase patients whose transfusions did not lead to a DHTR event (the ‘*VOC’* group), acute-phase patients whose transfusions led to a DHTR event (the ‘*pre-DHTR’* group) and patients from whom samples were collected during a post-transfusion hemolytic process (the ‘DHTR’ group) (**Figure 1B**). LIGHT levels were significantly lower in pre-DHTR and DHTR samples (mean values: 179.5 and 165.9 pg/mL, respectively) than in plasma samples from acute-phase patients who did not experience a DHTR (VOC; mean value: 419.9 pg/mL). LIGHT levels did not differ significantly between pre-DHTR and DHTR patient samples for either total DHTR (*n* = 21) or paired DHTR (pre-post patients, *n* = 11) events (**Figure 1C** and **Supplemental Fig. S1D**).

### Correlations of plasma LIGHT concentrations with Hb levels and proinflammatory factors

Previous data revealed that ELISA-determined LIGHT levels were related to LDH and white blood cell (WBC) counts, specifically in hydroxyurea (HU)-treated SCD patients^8^, but with no correlation detected with hemoglobin (Hb) levels, hematocrit (Hct), or reticulocyte counts. We investigated the relationship between LIGHT concentrations and hematological parameters or cytokine levels in our cohort as a function of the clinical state of the selected patients. The data compiled for twenty-eight ‘*Basal*’ and thirty-three ‘*Acute*’ SCD patients revealed a small but significant association with Hb levels in both patient groups (**Figures 1F and G**). Plasma bilirubin and LDH levels, and secretory phospholipase A_2_ (sPLA_2_) levels were determined by ELISA, when possible, but were insufficiently documented for these patients for the results to be exploitable (data not shown). We also assessed the relationships LIGHT levels in SCD plasma samples and pro-inflammatory factor levels. We found a small but significant association of LIGHT levels with IL-27 (*p* = 0.0147) and TNFα (*p* = 0.0242) levels in the plasma of basal SCD patients, and with interferon (IFN)γ (*p* = 0.0003) and TNFα (*p* = 0.0164) levels in the plasma of acute patients (**Figures 1F and G**). Conversely, no association was found with ICAM-1 in pre-DHTR and DHTR patients (**Supplemental Fig. S1F**), whereas a significant correlation was observed with macrophage inflammatory protein (MIP)-1β levels in DHTR patients (*p* = 0.0094; **Supplemental Fig. S1G**). All the relationship tested between LIGHT levels and the various pro-inflammatory factors quantified are shown in **Supplemental fig. S2**.

### Soluble human recombinant LIGHT activates endothelial cells *in vitro*

LIGHT is known to be a potent activator of ECs and could, therefore, contribute to endothelial activation in SCD^8^. We tested this hypothesis in our conditions, by assessing the potential effect of hrLIGHT, at various doses, on HUVECs cultured in plastic dishes (static conditions). We assessed the expression of endothelium activation markers by flow cytometry (**Figure 2A**) and fluorescence microscopy (**Figure 2B**). The expression of activation markers on EC membranes was assessed by flow cytometry, focusing on intercellular adhesion molecule 1 (ICAM-1/CD54), von Willebrand factor (vWF), E-selectin (CD62-E) and vascular cell adhesion molecule 1 (VCAM-1/ CD106). HUVECS were either non-treated (NT) or were treated for 4, 8 or 20 hours with LIGHT (50, 100 or 200 ng/mL). Treatment with soluble hrLIGHT induced a time dependent activation of HUVECs associated with the detection of E-selectin and VCAM-1/ CD106, beginning 4 h after the start of treatment, followed by ICAM-1 and vWF after 8 hours of treatment. We observed classic type II activation, at LIGHT doses of at least 50 ng/mL (**Figure 2A**). Nevertheless, no dose-dependence for hrLIGHT treatment was detected for these markers. HUVEC activation was also confirmed by fluorescence microscopy, notably with CD106/VCAM-1 and CD54/ICAM-1 (**Figure 2B**). Treatment with hrLIGHT did not modify the HUVEC monolayer, as attested by the conservation of CD31/PECAM-1 expression.

**Figure 2.**
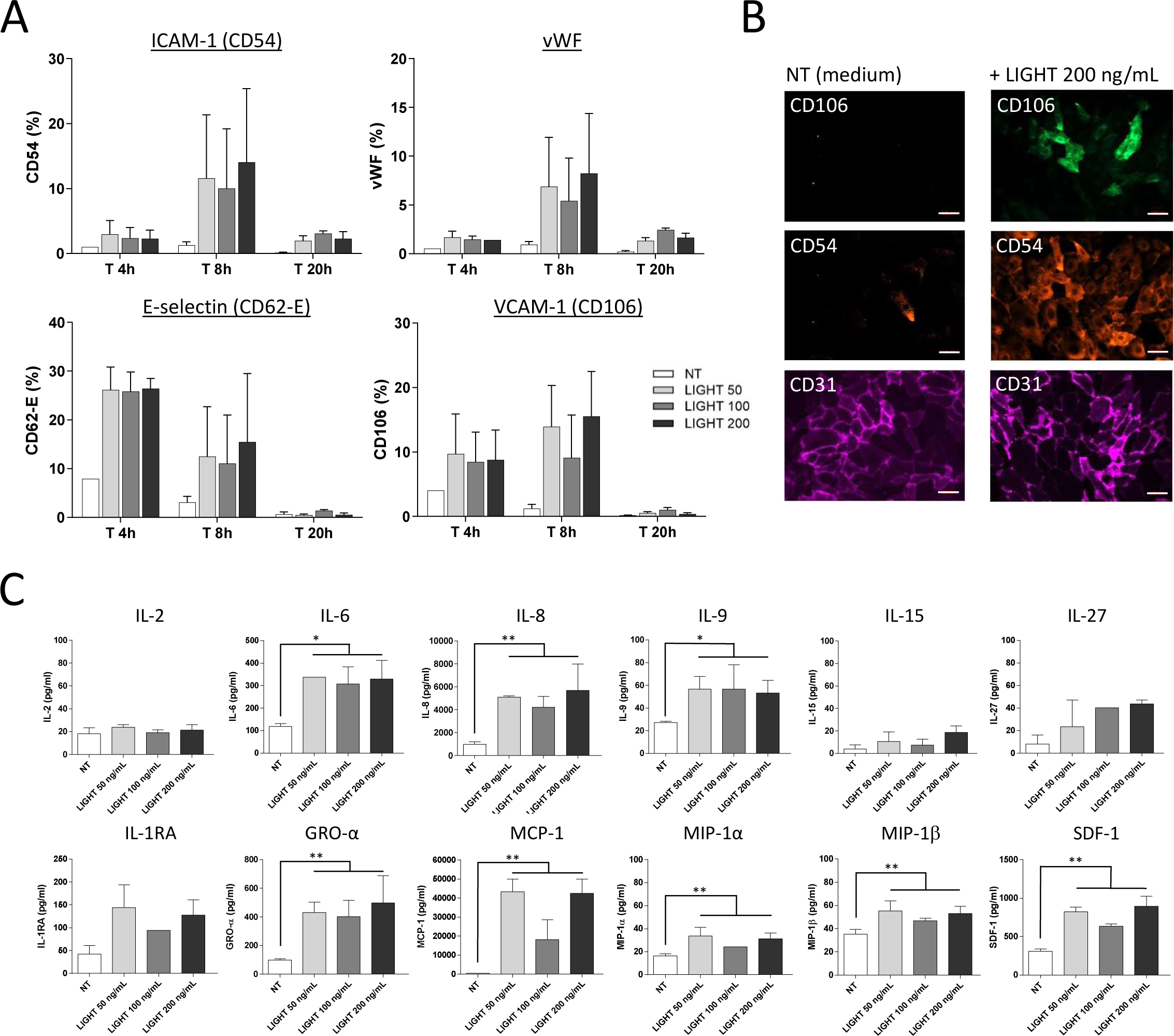
Activation of HUVECs after treatment (time- and dose-dependent responses) with recombinant LIGHT. **(A)** The effect of hrLIGHT (50, 100 and 200 ng/mL) on the expression of adhesion molecules on HUVECs after 4, 8 and 20 hours of treatment was assessed. Stimulated cells were washed and harvested with cold PBA/lidocaine with gentle shaking at +4 °C. Detached cells were then rinsed and stained by incubation for 15 minutes at +20 °C with the following monoclonal antibodies (mAbs): BB515-conjugated anti-CD54, PE-conjugated anti-vWF, APC-conjugated anti-CD106, BV421-conjugated E-selectin and BV510-conjugated anti-CD31. The cells were then fixed before analysis on a BD Canto II flow cytometer. **(B)** The HUVECs cultured in fibronectin-coated Ibidi µ-Slide I were left untreated or were treated with hrLIGHT (50, 100 and 200 ng/mL) for 4 hours at +37 °C. The cells were then washed and stained by incubation for 15 minutes at +20 °C with the following mAbs: BB515-conjugated anti-CD54, APC-conjugated anti-CD106 and BV510-conjugated anti-CD31. They were then washed with PBS and mounted in DAPI-containing medium. Horizontal scale bars represent 50 µm. Data were collected on a Axio Imager Zeiss Z1 (Carl Zeiss, Marly le Roi, France) microscope and analyzed with Zen software. **(C)** HUVECs were treated for 4 hours with hrLIGHT (50, 100 and 200 ng/mL) and cytokine release was then assessed. TNFα (20 ng/mL) was used as a positive control (not shown); mock conditions, identical except that hrLIGHT was not added, were tested in parallel (not shown on the graphs). Previously harvested culture supernatants were used for the quantification of the following cytokines: IL-2, IL-6, IL-8, IL-9, IL-15, IL-1RA, TNFα, GRO-α, MCP-1, MIP-1α, MIP-1β and SDF-1; each sample was tested in duplicate. Concentrations are shown as histograms +SEM error bars. *, ** indicate values of *p*<0.05, and 0.01, respectively, for comparisons of the LIGHT-treated group with unstimulated cells. The horizontal scale bars represent 50 µm.

### Recombinant LIGHT induces a proinflammatory secretory profile in cultured HUVECs

We then quantified 34 soluble activation factors potentially secreted into the HUVEC culture medium in response to hrLIGHT, by the Luminex technique (**Figure 2C**). The levels of several cytokines, including IL-6, IL-8 and IL-9, increased significantly after four hours of hrLIGHT treatment (with a similar but non-significant trend for IL-1RA and IL-27). Cytokine release was also significantly stimulated for MCP-1, MIP-1α, MIP-1β, GRO-α, and SDF-1 (ANOVA relative to untreated cells).

### HUVEC activation by LIGHT leads to physiological flow adherence of sickle (SS) platelets

In a closed flow culture *in vitro* assay using Ibidi µ-Slide devices coated with fibronectin and seeded with primary HUVECs investigated whether the activation by LIGHT treatment led to adherence of blood components (**Figures 3 and supplemental fig. S3, S4 and S5**). The AA blood samples were obtained from healthy donors; the SS blood samples were obtained from patients with steady-state SCD followed during consultations, outside of painful events. For the preconditioning step, HUVECs were either left untreated (‘*NT*’) or were treated with hrLIGHT (500 ng/mL), which was added to the culture medium for four hours under physiological flow conditions (**Figure 3A**). We found that four hours of a LIGHT treatment on biochips strongly activated ECs, as shown by their expression of CD54/ICAM-1 (**Figure 3B**, lower panel). When fresh SS blood samples from patients were tested in the functional flow assay on LIGHT-treated HUVECs, we observed strong adherence of SS platelets (visualized through the expression of CD41a/ platelet glycoprotein 2b) that was not observed for untreated HUVECs (**Figure 3B,** upper panel). With few exceptions, CD45^+^ leukocytes did not adhere to the biochip, whether AA or SS blood was used in the test. A quantification of adherent platelets showed there to be only few adherent platelets per field after flow period with AA whole blood, with or without LIGHT pretreatment. With the pathological SS blood, the mean values were much higher, with 60.9 ± 32 adherent platelets/mm^2^ for LIGHT-treated HUVECs (and 9.0 ± 4.8 platelets/mm^2^ for untreated cells). Such adherence was clearly observed with both LIGHT-activated ECs and the EC extracellular matrix on biochips (yellow arrowhead; **Figure 3B,** lower panel, on the right). During acute events in SCD patients, sickle (SS) RBCs display an increase in adherence to ECs and/or proteins of the vascular wall^25, 28^. Such abnormal interactions may play a crucial role in the occurrence of vaso-occlusive events in this disease. Then, we questioned if that the activation of ECs by LIGHT induced an increase in RBC adherence. Such treatment of the cells (right panel) led to adherence of many more SS RBCs than for untreated HUVECs (left panel) (**Supplemental fig S4 and S5**). The increase in the number of AA RBCs adhering after the LIGHT treatment of HUVECs was non-significant when compared to untreated HUVECs.

**Figure 3.**
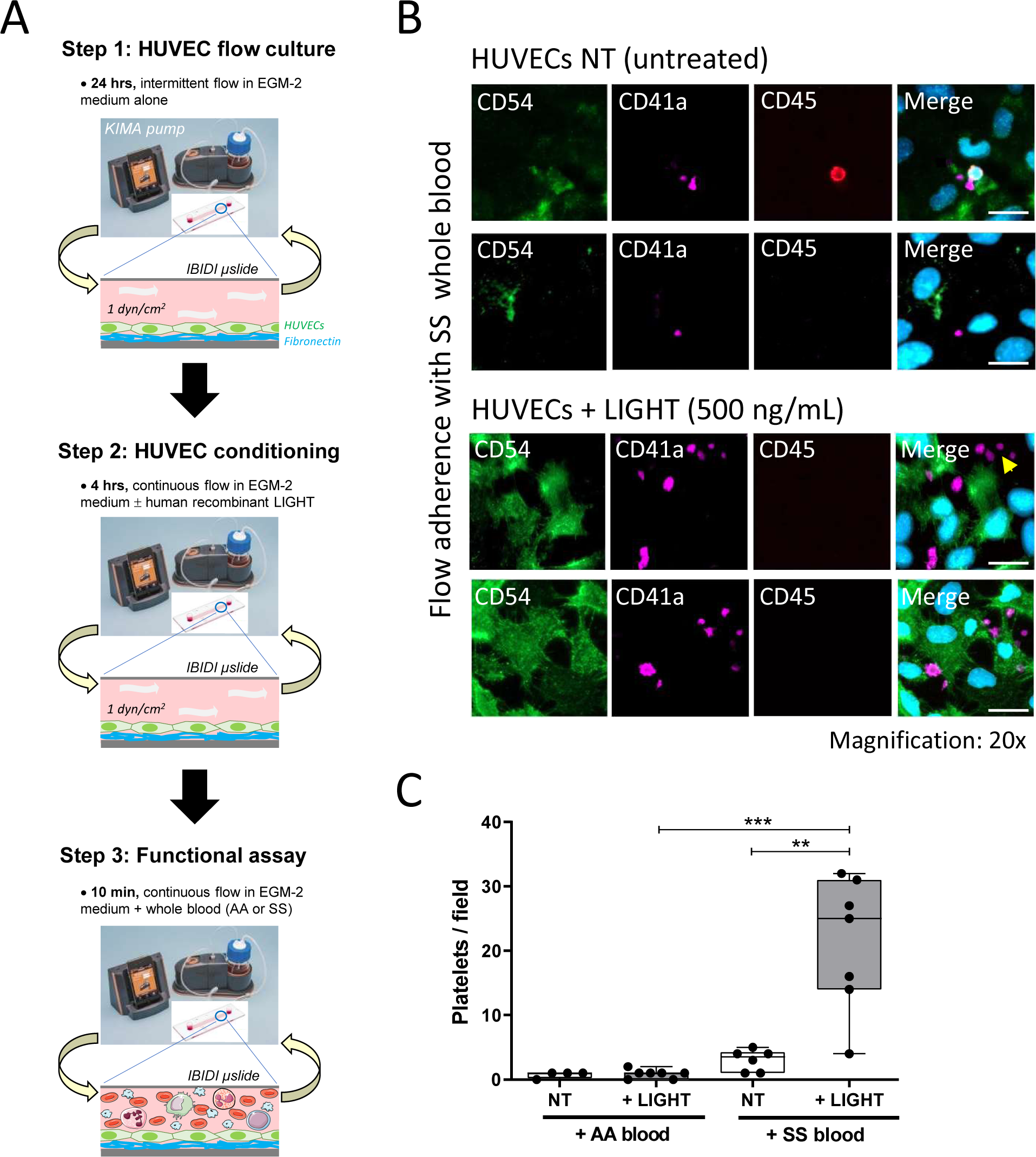
Flow adherence tests on whole blood from SS with LIGHT-activated HUVECs revealing adherence of sickle platelets. HUVECs in an Ibidi µ-Slide microfluidic device were conditioned by perfusing the microfluidic device at a flow rate of 350 µL/min (1 dyne/cm^2^ shear stress) for 4 hours. They were left untreated or were treated with hrLIGHT (500 ng/mL) in EGM-2 medium, which was infused through the system. The system was then perfused with fresh whole blood from AA donors or SS patients for 10 min. Washing with EGM-2 was then performed to remove all circulating blood cells. The cells remaining in the Ibidi µ-Slide device were stained by incubation for 15 minutes at +20 °C with the following mAbs: BB510-conjugated anti-CD54, BB515-conjugated anti-CD41a or APC-conjugated anti-CD45. They were then mounted in DAPI-containing medium. Data were collected on a Axio Imager Z1 with Zen software. The yellow arrowhead indicates platelet adherence to an EC-free extracellular matrix zone. The horizontal scale bars represent 50 µm.

### Neutralization both HUVEC activation and sickle (SS) platelet adherence by anti-LIGHT Abs

Finally, the results we obtained concerning a possible reversion effect after LIGHT blockade showed that increasing doses of neutralizing polyclonal anti-LIGHT Ab induced a dose-dependent reduction of the expression of HUVEC activation markers **(Figure 4A and B**), and of the adhesion of sickle (SS) platelets in physiological flow experiments **(Figure 4C**). A decrease in the activation state of the cells was confirmed by the levels of the three membrane markers detected (CD54, CD106 and vWF; **Figure 4A and B**), and by the release of various pro inflammatory cytokines (**Figure 4D**). For the adherence of sickle (SS) platelets in physiological flow experiments, no decrease was observed with a control non-specific IgG, supporting a role for LIGHT in the first step in vaso-occlusion in SCD.

**Figure 4:**
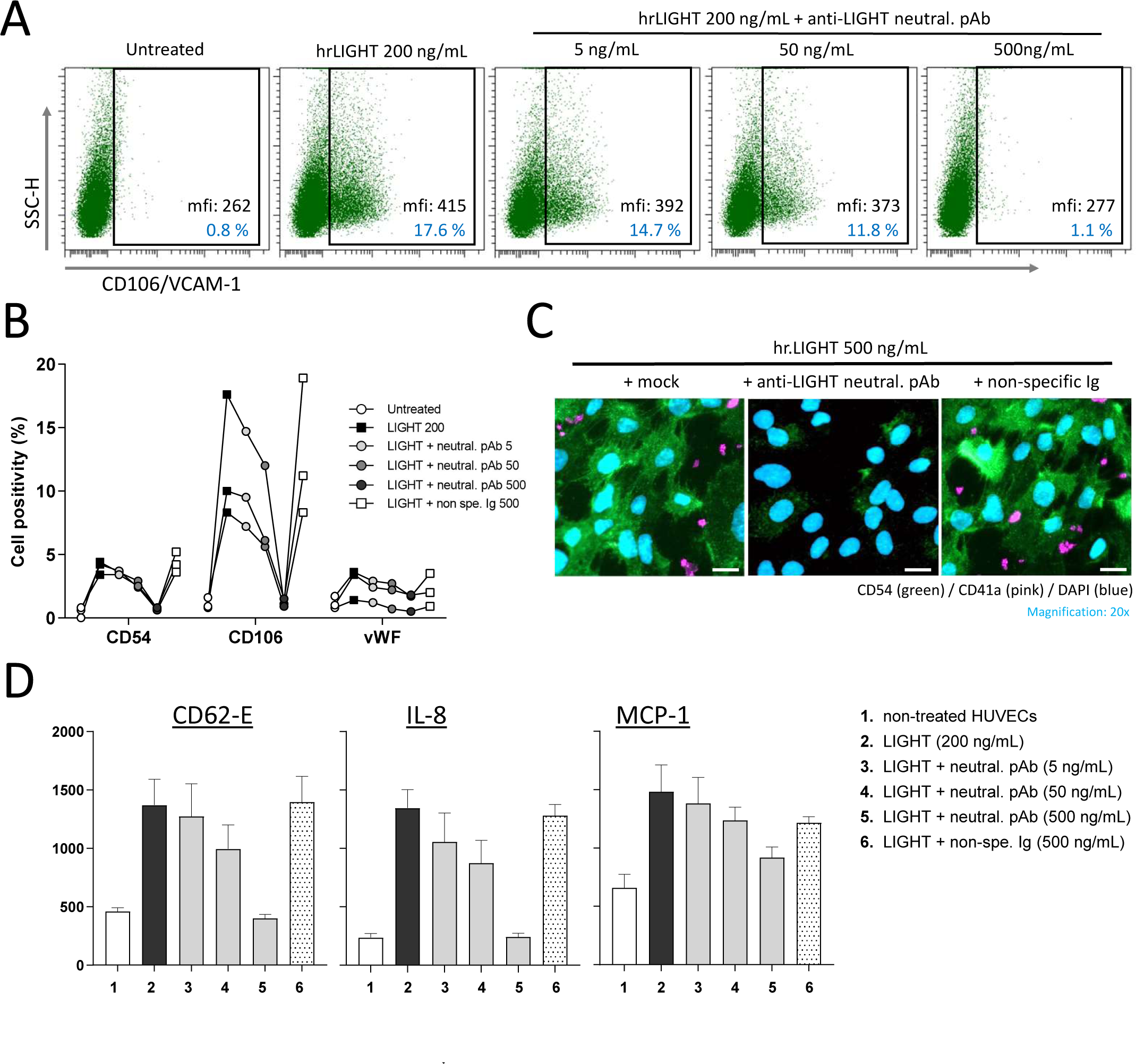
Blocking effect of an anti-LIGHT antibody on HUVEC activation and sickle (SS) platelet adherence. **(A)** The effects on the expression of adhesion molecules on HUVECs of hrLIGHT (200 ng/mL), a rabbit polyclonal blocking Ab directed against LIGHT and a nonspecific rabbit Ig were tested after 6 hours of treatment. Cells were harvested in cold PBA/lidocaine, rinsed and then stained with anti-CD54, anti vWF, anti-CD106, anti-E-selectin and anti-CD31 mAbs for analysis on a BD Canto II flow cytometer. In **(B)** the results obtained for CD106/VCAM-1 are shown for the various sets of conditions: untreated HUVECs (white circles), 200 ng/mL LIGHT alone (black squares), 200 ng/mL LIGHT + 5 ng/mL anti-LIGHT neutralizing Abs (light gray circles), or + 50 ng/mL anti-LIGHT neutralizing Abs (gray circles) or + 500 ng/mL anti-LIGHT neutralizing Abs (dark gray circles) and, finally, 200 ng/mL LIGHT + 500 ng/mL non-specific Ig (black circles) **(C)** HUVECs in the Ibidi µ-slide device, with or without hrLIGHT (500 ng/mL) treatment, were conditioned at a flow rate of 350 µL/min for 4 hours. The system was then perfused for 10 minutes with fresh whole blood from an SS patient before being washed. Adherent cells were stained with BB510-conjugated anti-CD54 and BB515-conjugated anti-CD41a Abs, mounted in DAPI-containing mounting medium and analyzed on an Axio Imager Z1 with Zen software.

**Figure 5.**
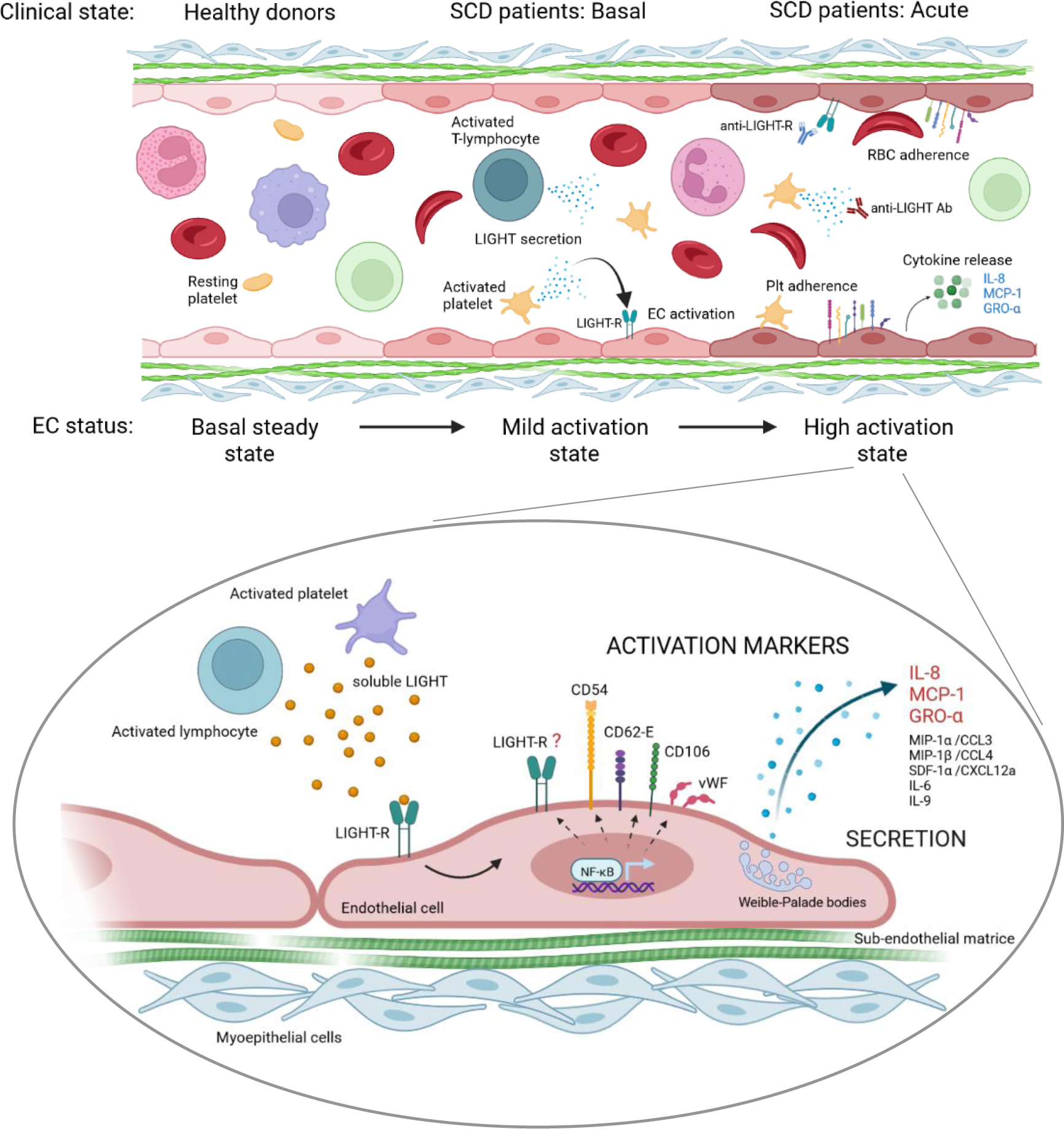
Schematic abstract, model for HUVEC activation by soluble LIGHT.

## DISCUSSION

Soluble LIGHT is already known to be significantly elevated in the plasma of SCD patients, compared to healthy volunteers. It represents now a novel immune checkpoint molecule that plays essential roles in both innate and acquired immunity. In the present study, we extend previous finding by showing that LIGHT levels are significantly increased in SCD plasmas from patients during the acute phases of the disease like painful occlusive crises events (VOC) associated or not with ACS, with a 3-fold increasing factor compared to healthy donors, a result which is consistent with previous data from Garrido and colleagues^8^.

Comparisons made between the clinical phases of SCD were allowed with our cohort, where patients were included for a reason of transfusion either during a basal steady-state or an acute phase. In the latter acute cases, LIGHT levels in plasma were markedly higher than in the ‘*Basal*’ state counterparts, a result never documented before. We could conclude that LIGHT levels are linked to clinical severity of the disease in homozygous SCD adult patients. In adult SCD patients, our data and results of other teams^18, 19, 29^ observed a link between LIGHT levels and plasma LDH (mainly for HU-treated patients) but also with CD40-L, IL8 and CD54/ICAM1 both upregulated in ECs after a LIGHT stimulation; others detected an upregulation of the Tissue Factor^10^. It is of interest to notice that the upregulation of CD54 we detect reproduces the effect of platelet-derived CD40L on endothelial cells, another member of the TNF superfamily^30^.

HU treatment improves clinical manifestations in SCD and can modify various hematological blood parameters^31, 32^. In our cohort, we observe a trend but significant correlation link between soluble plasma LIGHT and Hb levels but also with some inflammatory markers in SCD patients depending on the clinical phase considered with to particularly notice among them IFNγ in acute painful patients (R^2^ = 0.4791; p = 0.0003). In our selected cases of SCD adult patients, no effect of a HU-treatment was detected concerning correlations of LIGHT with hematological parameters. Finally, we tested if LIGHT was correlated with sPLA_2_, an enzyme that is secreted during the acute phases in SCD and specially during ACS^33, 34^; sPLA, by generating lysophosphatidic acid, may result in vascular dysfunction. No correlation between plasma sPLA_2_ and LIGHT was observed in our cohort (data not shown). Concerning LDH, only few data collected from the patient’ cohort and were not sufficient to test or confirm a correlation link with LIGHT levels^8^.

In SCD, the oxidative stress first, leads to activation of ECs in the vessel walls and contributes to increase the adherence of cell blood components in the vessel walls. The sickle RBCs then abnormally adhere to the vascular endothelium, contributing to microvascular occlusions, driving hospitalizations^39, 41^. Pro-inflammatory proteins in the circulation plays also an important role in the activation of both the endothelium and leucocytes^35^. LIGHT/TNFSF14 pro-inflammatory, pro-thrombotic and atherogenic cytokine could be now considered as a high effective activating circulating factor of ECs^10, 18, 36^, an activation we confirmed under static conditions. The fact that a one and only soluble factor, the LIGHT cytokine, detected high in the circulation of SCD patients, could directly participate to endothelium activation, then initiating the vascular inflammation associated with the pathophysiology of SCD, might be of particular interest for the management of the disease.

Platelets can adhere to activated endothelium^37, 38^ and this initiate and sustain inflammatory reactions at this site. Adherence of SS RBCs can be explained by the network of adhesion molecules and the interactions between them: the Lutheran/basal cell adhesion molecule (Lu/BCAM) on endothelium with the α4β1 integrin on SS reticulocytes, first; then Lu/BCAM with the intercellular adhesion molecule 4 (ICAM-4) on SS RBCs^39, 40^.Through physiologic flow experiments into biochips, we provide the first evidence for a direct link between the LIGHT-induced endothelial activation and adherence of blood cell components. Then, the LIGHT-induced secondary-step adherence of both sickle red cells and platelets following EC’ activation can potentially be the first step in vaso-occlusions^41^. Such events have been shown to initiate vaso-occlusive mechanisms in animal models^35^ as well as in SCD patients^42, 43^. Finally, the impact of the relative density of sickle SS RBC populations injected in microfluidic experiments, a main theme of study regarding the SCD physiopathology in our group^44–45^, was not tested in our conditions.

As a limitation of the study, it may be argued that the rhLIGHT concentrations tested in the *in vitro* experiments in the present study were too high. If the values obtained from 50 - 200 ng/mL treatments were sufficient to induce a high HUVEC’ activation state under static conditions *in vitro*, flow experiment assays necessitated higher amounts of LIGHT to observe any reproductible effect. This may probably be due to unwanted deposit of LIGHT molecules onto plastic sides into the microfluidic system by adsorbance (probably in the flexible tubes and/or inside the pump mechanism).

As a conclusion, soluble LIGHT directly contributes to endothelial inflammation in the context of SCD, with this soluble factor elevated in most of the patients we tested, in association with other markers of endothelial activation. Better understanding the link between LIGHT-induced EC’ activation and SCD pathophysiology could help in treating complications of SCD. The LIGHT-driven inflammation could represent a pathogenic loop in SCD, representing a potential target by using blocking anti-LIGHT receptor on ECs or anti-soluble LIGHT neutralizing antibodies^19^. Very recently, blockade of LIGHT with recombinant HVEM-Fc fusion protein recently showed to ameliorate liver injury in poly(I:C)-induced hepatitis^46^; a human LIGHT-neutralizing antibody, CERC-002 might reduce the risk of respiratory failure and death in COVID-19–related cytokine release and acute respiratory distress syndromes^47, 48^. Future investigations must be conducted to determine the extent of the contribution of LIGHT to SCD clinical complications, and whether it could represent a potential novel therapeutic target cytokine.

## ABREVIATIONS

TNFSF14: tumor necrosis factor superfamily member 14
HUVECs: human umbilical vein endothelial cells
SCD: sickle-cell disease
RBCs: red blood cells
ECs: endothelial cells
ICAM-1: intercellular adhesion molecule-1
VCAM-1: vascular cell adhesion molecule-1
vWF: von willebrand factor
VOC: vaso-occlusive crisis
ACS: acute chest syndrome
DHTR: delayed hemolytic transfusion reaction
LDH: lactate dehydrogenase
Hb: hemoglobin
Hct: hematocrit
sPLA2: secreted phospholipase A2.

## Acknowledgements

The authors thank laboratoire d’excellence GR-Ex, Paris, France. We are grateful to the patients and the healthy blood donors who participated in this study, and the *Etablissement Français du Sang* laboratories responsible for collecting blood donations. We very thank Dr Benoît Vingert for constructive scientific discussions and technical assistance on Luminex.

## Author contributions

PC was the principal investigator and take primary responsibility for this article. GB, KN-P, GDL and PC performed the laboratory work for the study. GB and PC collected and analyzed the data and performed the statistical analysis. PC wrote the manuscript. FP and PB reviewed the manuscript. PC, FP and PB coordinated the research. All the authors contributed to the manuscript and approved the final version of the manuscript.

## SUPPLEMENTAL MATERIAL

### SUPPLEMENTARY METHODS

#### SCD patient samples, RBC collection and preparation

A cohort of adult homozygous HbS patients (age ≥ 18 years) with a definitive diagnosis of SCD (27 men/34 women; mean age: 34.7 ± 8.5 years) was studied during follow-up for transfusion, after admission to the Henri Mondor Hospital (Créteil, France), or during their follow-up in the Sickle Cell Referral Center. We prospectively collected pre- and post-transfusion blood samples from all the SCD patients^22, 26^ (*n* = 108, with some included in more than one group), and these samples were then assigned to the various categories. Acute-phase patients whose transfusion did not lead to hemolytic transfusion reactions were assigned to the ‘*VOC*’ group. Finally, a group of 21 adult healthy volunteers of African descent (*HD*; 10 men/11 women; mean age: 30.9 ± 10.1 years) were selected from the *Etablissement Français du Sang* cohort of blood donors. Written informed consent was obtained from all patients and donors.

#### Luminex detection assays for cytokines and inflammatory markers

Plasma samples from the cohort and culture supernatants were thawed at +4°C and tested in a multiplex cytokine assay performed with Luminex technology (Affymetrix, Vienna, Austria)^22^. Plasma samples were first centrifuged for 10 min at 10,000 x *g*, +4 °C, to remove any remaining lipids and platelets. We tested a panel of 12 selected cytokines, chemokines, and growth factors, as follows, to determine the activation profile of HUVECs: interleukin (IL)-2, IL-6, IL-8, IL-9, IL-15, IL-1RA, TNFα, GRO-α, MCP-1, MIP-1α, MIP-1β and SDF-1. Each sample was tested in duplicate and at two dilutions (pure and 1:10 dilution) for each time point. Data were collected on a MAG*PIX* (XMAP Technology) and analyzed with FlowCytomics 2.4 software (Affymetrix).

#### Primary HUVECs: isolation and culture

HUVECs were isolated from fresh umbilical cord veins obtained from the gynecology-obstetrics department of the *Centre Hospitalier Intercommunal de Créteil* (CHIC, Val-de-Marne). Umbilical cords were washed with saline and phosphate buffered saline (PBS; Gibco, Life Technologies, Courtabœuf, France) to remove all traces of blood. They were cut at both ends and the umbilical vein of the cord was rinsed with PBS via a cannula and syringe. Primary ECs were detached by incubation with 0.5 mg/mL collagenase (Sigma Aldrich) at +37 °C for 10 min and the resulting cell suspension washed with endothelial growth medium-2 (EGM-2) complete medium (Lonza LTD, Basel, Switzerland) supplemented with 20 % fetal bovine serum (PAA Laboratories, Velizy Villacoublay, France). After centrifugation (10 min at 800 x *g*), the cell pellet was resuspended in EGM-2 complete medium supplemented with 10 % fetal bovine serum. The HUVECs were cultured and amplified *in vitro* in BD Falcon plastic flasks (BD Life Sciences, Le-Pont-de-Claix, France) coated with 0.5 % gelatin (Sigma Aldrich) at +37 °C, under an atmosphere containing 5 % CO_2_ in a humidified incubator. In all experiments, the HUVECs were used at their fourth to sixth passage; their phenotype was regularly checked by CD31/PECAM-1 staining and analyse by flow cytometry.

## SUPPLEMENTAL FIGURES

Supplemental fig. S1. **Measurements of LIGHT/TNFSF14 levels in plasma samples from SCD patients and healthy donors.**

**(A)** Pretransfusion plasma samples from SCD patients in a basal steady state (*n* = 82; gray circles) and a control group of blood donors (*n* = 21; white circles) were subjected to ELISA for the quantification of soluble LIGHT. **(B)** ROC curve analysis of the LIGHT levels in SCD plasma samples relative to donor samples; AUC: area under the curve. **(C)** Pretransfusion plasma samples from SCD patients in a basal steady state (Basal; *n* = 28; white squares) or in a painful acute phase (Acute; *n* = 33; black squares) were tested by ELISA to quantify soluble LIGHT factor. In **(D)**, the LIGHT levels in SCD plasma samples were also compared before and at the onset of a DHTR; *ns*: non-significant. **(E)** A significant correlation was found between plasma LIGHT concentration and plasma levels of Hb levels, IL-27 and IL-6 in pretransfusion samples from the whole population of SCD patients (‘Basal’: white squares; ‘Acute’: black squares; ‘pre-DHTR’: white circles). **(F)** The significant correlation between Hb levels and LIGHT concentrations in plasma samples from the ‘pre-DHTR’ and ‘DHTR’ categories (left and right, respectively) of SCD patients is shown. **(G)** The significant correlations between plasma LIGHT concentration and ICAM-1 and MIP-1β levels in plasma samples from the ‘pre-DHTR’ and ‘DHTR’ categories (left and right, respectively) of SCD patients are represented. Values of *p* are indicated on the graphs (Mann-Whitney *U* test).

Supplemental fig. S2. **Correlations between the levels of various cytokines and soluble LIGHT in plasma samples from the ‘Basal’ and ‘Acute’ groups of SCD patients.**

The correlations obtained by analyses of linear regression between plasma LIGHT levels and the levels of the following cytokines are shown: IL-6, IL-10, IL-27, IFN-γ, TNFα, ICAM-1, IP-10, MIP-1β and RANTES, for plasma samples collected from ‘Basal’ (upper panel) and ‘Acute’ (lower panel) SCD patients.

Supplemental fig. S3. **Flow experiments with HUVECs activated by recombinant LIGHT, leading to flow adherence of sickle (SS) platelets** (magnification of the images in Figure 3).

HUVECs in an Ibidi microfluidic device were conditioned by perfusing the microfluidic device at a flow rate of 350 µL/min (1 dyne/cm^2^ shear stress) for 4 hours. They were left untreated or were treated with hrLIGHT (500 ng/mL) in EGM-2 medium, which was infused through the system. Finally, the system was perfused with whole blood from an AA donor or SS patient for 10 minutes. It was then washed to remove all circulating blood cells. The cells in the Ibidi were stained by incubation for 15 minutes at +20 °C with the following antibodies: BB510-conjugated anti-CD54, BB515-conjugated anti-CD41a, or APC-conjugated anti-CD45 antibodies. They were then mounted in DAPI-containing medium. The horizontal scale bars represent 50 µm. Data were collected on an Axio Imager Z1 equipped with Zen software. The horizontal scale bars represent 50 µm.

Supplemental fig. S4. **Flow adhesion of red blood cells to HUVECs with and without LIGHT conditioning.** The HUVECs in an Ibidi µ-Slide were subjected to intermittent flow overnight with the KIMA pump system and were then conditioned by perfusing the microfluidic device at a flow rate of 350 µL/min (1 dyne/cm^2^ shear stress) for 4 hours. They were left untreated or were treated with hrLIGHT (500 ng/mL) in EGM-2 medium, which was infused through the system. The system was perfused with RBCs isolated from fresh blood samples, washed in HBSS and diluted in EGM-2. Adhesion was measured under flow conditions with a Mirus Nanopump and quantified by phase-contrast microscopy on an Axio Imager Z1 (Carl Zeiss, Marly le Roi, France) with a black and white Axiocam 702 camera and VenaFlux Assay 2.3 analysis software (Cellix). Adherent RBCs were counted by a person blind to the patient’s clinical status and treatments, in five representative areas along the centerline of the biochip for each set of conditions.

Supplemental fig. S5. **Flow experiments with HUVECs activated by recombinant LIGHT, leading to adherence of sickle (SS) platelets** (magnification of the images in Supplemental fig. S4).

Supplemental fig. S6. **Adherence of sickle platelets to HUVECs activated by recombinant LIGHT is abolished by treatment with anti-LIGHT antibody** (magnification of the images in Figure 4). HUVECs seeded in fibronectin-coated Ibidi µ-Slide device for fluidic experiments were treated with recombinant hrLIGHT (500 ng/mL) alone, with hrLIGHT 500 ng/mL + anti-LIGHT neutralizing pAb 500 ng/mL or with hrLIGHT 500 ng/mL + non-specific rabbit Ig 500 ng/mL and conditioned at a flow rate of 350 µL/min for 4 hours. Then SS patient’ fresh whole blood was perfused for 10 min, followed by a washing step. Adhered cells were stained with BB510-conjugated anti-CD54 and BB515-conjugated anti-CD41a, mounted with a DAPI-containing medium and analyzed on a Axio Imager Z1 equipped with Zen software. The horizontal scale bars represent 50 µm

## References

1. Kato GJ, Piel FB, Reid CD, et al. Sickle cell disease. Nat Rev Dis Primers 2018;418010

2. Ataga KI, Kutlar A, Kanter J, et al. Crizanlizumab for the prevention of pain crises in sickle cell disease. N Engl J Med 2017;376(5):429–439

3. Adams RJ, McKie VC, Hsu L, et al. Prevention of a first stroke by transfusions in children with sickle cell anemia and abnormal results on transcranial Doppler ultrasonography. N Engl J Med 1998;339(1):5–11

4. Ballas SK, Lusardi M. Hospital readmission for adult acute sickle cell painful episodes: frequency, etiology, and prognostic significance. Am J Hematol. 2005;79(1):17–25.

5. Vichinsky EP, Neumayr LD, Earles AN, et al. Causes and outcomes of the acute chest syndrome in sickle cell disease. National Acute Chest Syndrome Study Group. N Engl J Med 2000;342(25):1855–1865

6. Bartolucci P, Habibi A, Khellaf M, et al. Score predicting acute chest syndrome during vaso-occlusive crises in adult sickle-cell disease patients. EBioMedicine 2016;10305–311

7. Ballas SK, Larner J, Smith ED, Surrey S, Schwartz E, Rappaport EF. Rheologic predictors of the severity of the painful sickle cell crisis. Blood. 1988;72(4):1216–23

8. Garrido VT, Proença-Ferreira R, Dominical VM, et al. Elevated plasma levels and platelet-associated expression of the pro-thrombotic and pro-inflammatory protein, TNFSF14 (LIGHT), in sickle cell disease. Br J Haematol. 2012;158(6):788–97.

9. Mauri DN, Ebner R, Montgomery RI, et al. LIGHT, a new member of the TNF superfamily, and lymphotoxin alpha are ligands for herpesvirus entry mediator. Immunity. 1998;8(1):21–30.

10. Celik S, Langer H, Stellos K, et al. Platelet-associated LIGHT (TNFSF14) mediates adhesion of platelets to human vascular endothelium. Thromb Haemost. 2007;98(4):798–805.

11. Otterdal K, Smith C, Oie E, et al. Platelet-derived LIGHT induces inflammatory responses in endothelial cells and monocytes. Blood. 2006;108(3):928–35.

12. Petreaca ML, Yao M, Ware C, Martins-Green MM. Vascular endothelial growth factor promotes macrophage apoptosis through stimulation of tumor necrosis factor superfamily member 14 (TNFSF14/LIGHT). Wound Repair Regen. 2008;16(5):602–14.

13. Celik S, Shankar V, Richter A, et al. Proinflammatory and prothrombotic effects on human vascular endothelial cells of immune-cell-derived LIGHT. Eur J Med Res. 2009;14(4):147–56.

14. Holmes TD, Wilson EB, Black EV, et al. Licensed human natural killer cells aid dendritic cell maturation via TNFSF14/LIGHT. Proc Natl Acad Sci U S A. 2014;111(52):E5688–96.

15. Yan L, Da Silva DM, Verma B, et al. Forced LIGHT expression in prostate tumors overcomes Treg mediated immunosuppression and synergizes with a prostate tumor therapeutic vaccine by recruiting effector T lymphocytes. Prostate. 2015;75(3):280–91.

16. Xu G, Liu D, Okwor I, et al. LIGHT is critical for IL-12 production by dendritic cells, optimal CD4+ Th1 cell response, and resistance to Leishmania major. J Immunol. 2007;179(10):6901–9.

17. Dahl CP, Gullestad L, Fevang B, et al. Increased expression of LIGHT/TNFSF14 and its receptors in experimental and clinical heart failure. Eur J Heart Fail. 2008;10(4):352–9.

18. Sandberg WJ, Halvorsen B, Yndestad A, et al. Inflammatory interaction between LIGHT and proteinase-activated receptor-2 in endothelial cells: potential role in atherogenesis. Circ Res. 2009;104(1):60–8.

19. Brunetti G, Rizzi R, Storlino G, et al. LIGHT/TNFSF14 as a New Biomarker of Bone Disease in Multiple Myeloma Patients Experiencing Therapeutic Regimens. Front Immunol. 2018;23(9):2459

20. Skeate JG, Otsmaa ME, Prins R, et al. TNFSF14: LIGHTing the Way for Effective Cancer Immunotherapy. Front Immunol. 2020;11:922. Review

21. Madge LA, Kluger MS, Orange JS, May MJ. Lymphotoxin-alpha 1 beta 2 and LIGHT induce classical and noncanonical NF-kappa B-dependent proinflammatory gene expression in vascular endothelial cells. J Immunol. 2008;180(5):3467–77.

22. Mou Y, Yue Z, Zhang H, et al. High quality in vitro expansion of human endothelial progenitor cells of human umbilical vein origin. Int J Med Sci. 2017;14(3):294–301.

23. Chadebech P, de Ménorval MA, Bodivit G, et al. Cytokine changes in sickle-cell disease patients as markers predictive of the onset of delayed hemolytic transfusion reactions. Cytokine. 2020;136:155259.

24. Bencheikh L, Nguyen KA, Chadebech P, et al. Preclinical evaluation of the preservation of red blood cell concentrates by hypoxic storage technology for transfusion in sickle cell disease. Haematologica. 2022;107(8):1944–9.

25. Bartolucci P, Chaar V, Picot J, et al. Decreased sickle red blood cell adhesion to laminin by hydroxyurea is associated with inhibition of Lu/BCAM protein phosphorylation. Blood. 2010;116(12):2152–9

26. Chadebech P, Bodivit G, Di Liberto G, et al. Ex vivo activation of red blood cell senescence by plasma from sickle-cell disease patients: correlation between markers and adhesion consequences during acute disease events. Biomolecules. 2021;11(7):963.

27. Narbey D, Habibi A, Chadebech P, et al. Incidence and predictive score for delayed hemolytic transfusion reaction in adult patients with sickle cell disease. Am J Hematol. 2017;92(12):1340–1348.

28. Chaar V, Picot J, Renaud O, et al. Aggregation of mononuclear and red blood cells through an α4β1-Lu/basal cell adhesion molecule interaction in sickle cell disease. Haematologica. 2010;95(11):1841–8.

29. Chang YH, Hsieh SL, Chao Y, et al. Proinflammatory effects of LIGHT through HVEM and LTbetaR interactions in cultured human umbilical vein endothelial cells. J Biomed Sci. 2005;12(2):363–75.

30. Henn V, Steinbach S, Büchner K, et al. The inflammatory action of CD40 ligand (CD154) expressed on activated human platelets is temporally limited by coexpressed CD40. Blood. 2001;98(4):1047–54.

31. Roumenina LT, Chadebech P, Bodivit G, et al. Complement activation in sickle cell disease: Dependence on cell density, hemolysis, and modulation by hydroxyurea therapy. Am J Hematol. 2020;95(5):456–464.

32. Hebert N, Rakotoson MG, Bodivit G, et al. Individual red blood cell fetal hemoglobin quantification allows to determine protective thresholds in sickle cell disease. Am J Hematol. 2020;95(11):1235–1245.

33. Bargoma EM, Mitsuyoshi JK, Larkin SK, et al. Serum C-reactive protein parallels secretory phospholipase A2 in sickle cell disease patients with vasoocclusive crisis or acute chest syndrome. Blood. 2005;105(8):3384–5.

34. Styles LA, Abbou M, Larkin S, et al. Transfusion prevents acute chest syndrome predicted by elevated secretory phospholipase A2. Br J Haematol. 2007;136(2):343–4

35. Turhan A, Weiss LA, Mohandas N, et al. Primary role for adherent leukocytes in sickle cell vascular occlusion: a new paradigm. Proc Natl Acad Sci U S A. 2002;99(5):3047–51.

36. Halvorsen B, Santilli F, Scholz H, et al. LIGHT/TNFSF14 is increased in patients with type 2 diabetes mellitus and promotes islet cell dysfunction and endothelial cell inflammation in vitro. Diabetologia. 2016;59(10):2134–44.

37. Gawaz M, Neumann FJ, Dickfeld T, et al. Activated platelets induce monocyte chemotactic protein-1 secretion and surface expression of intercellular adhesion molecule-1 on endothelial cells. Circulation. 1998;98(12):1164–71.

38. Sachais BS. Platelet-endothelial interactions in atherosclerosis. Curr Atheroscler Rep. 2001;3(5):412–6

39. El Nemer W, Wautier MP, Rahuel C, et al. Endothelial Lu/BCAM glycoproteins are novel ligands for red blood cell alpha4beta1 integrin: role in adhesion of sickle red blood cells to endothelial cells. Blood. 2007;109(8):3544–51.

40. Chaar V, Laurance S, Lapoumeroulie C, et al. Hydroxycarbamide decreases sickle reticulocyte adhesion to resting endothelium by inhibiting endothelial lutheran/basal cell adhesion molecule (Lu/BCAM) through phosphodiesterase 4A activation. J Biol Chem. 2014;289(16):11512–21.

41. Hebbel RP, Yamada O, Moldow CF, et al. Abnormal adherence of sickle erythrocytes to cultured vascular endothelium: possible mechanism for microvascular occlusion in sickle cell disease. J Clin Invest. 1980;65(1):154–60.

42. Lanaro C, Franco-Penteado CF, Albuqueque DM, et al. Altered levels of cytokines and inflammatory mediators in plasma and leukocytes of sickle cell anemia patients and effects of hydroxyurea therapy. J Leukoc Biol. 2009;85(2):235–42.

43. Musa BO, Onyemelukwe GC, Hambolu JO, et al. Pattern of serum cytokine expression and T-cell subsets in sickle cell disease patients in vaso-occlusive crisis. Clin Vaccine Immunol. 2010;17(4):602–8.

44. Bartolucci P, Brugnara C, Teixeira-Pinto A, et al. Erythrocyte density in sickle cell syndromes is associated with specific clinical manifestations and hemolysis. Blood. 2012;120(15):3136–41.

45. Di Liberto G, Kiger L, Marden MC, et al. Dense red blood cell and oxygen desaturation in sickle-cell disease. Am J Hematol. 2016;91(10):1008–13.

46. Lai D, Lv Z, Lu X, et al. LIGHT amplification by NF-*κ*B contributes to TLR3 signaling pathway-induced acute hepatitis. Mediators Inflamm. 2023;2023:3732315.

47. Perlin DS, Neil GA, Anderson C, et al. Randomized, double-blind, controlled trial of human anti-LIGHT monoclonal antibody in COVID-19 acute respiratory distress syndrome. J Clin Invest. 2022;132(3):e153173.

48. Ware CF, Croft M, Neil GA. Realigning the LIGHT signalling network to control dysregulated inflammation. J Exp Med. 2022;219(7):e20220236.

