## Supplementary figures and images for "Soluble LIGHT (TNFSF14) activates endothelial cells, thereby priming the first vessel-occlusive events in acute sickle cell disease"

### Supplemental fig. S1

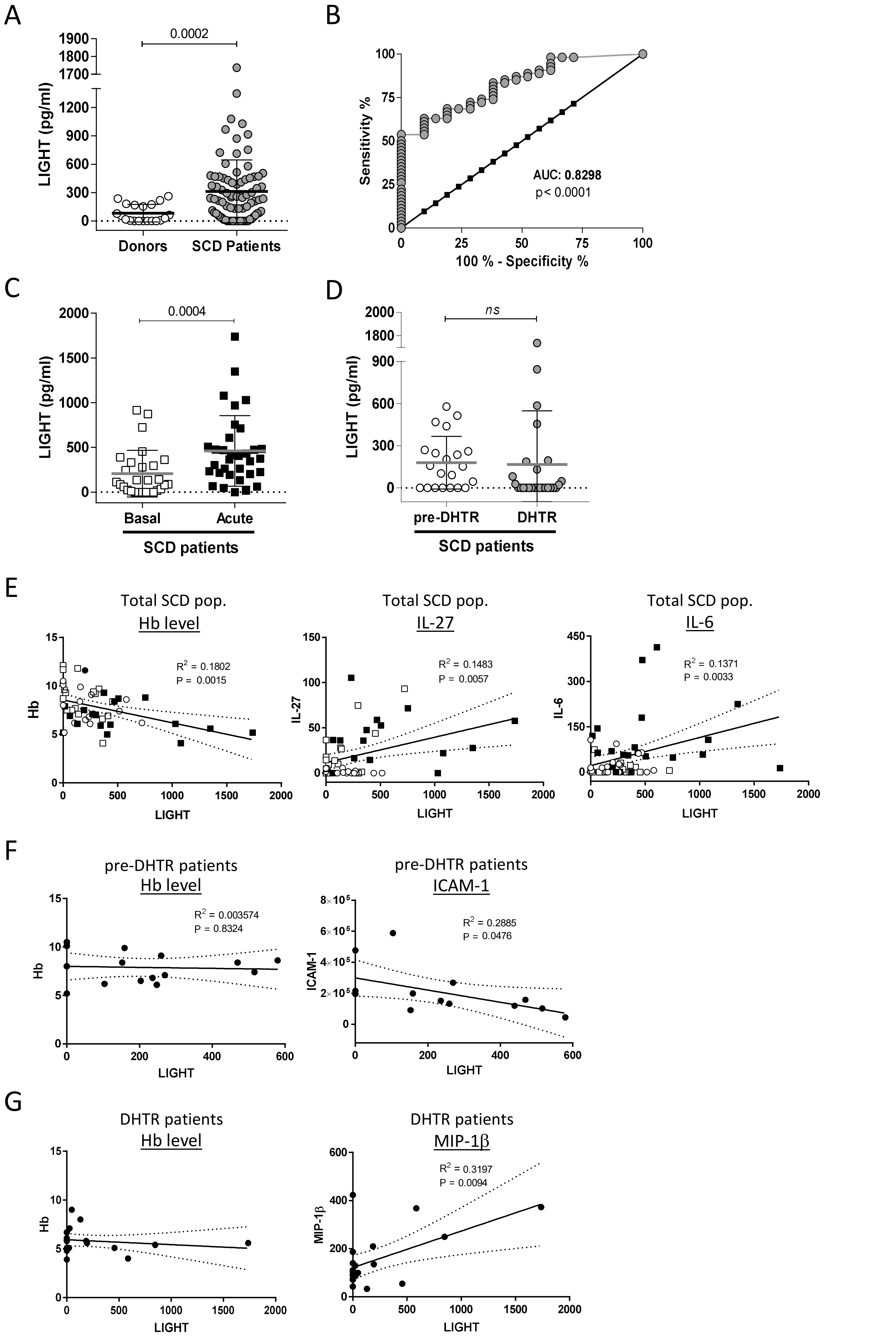


Supplemental figure S1

### Supplemental fig. S2

#### ‘Basal’ patients

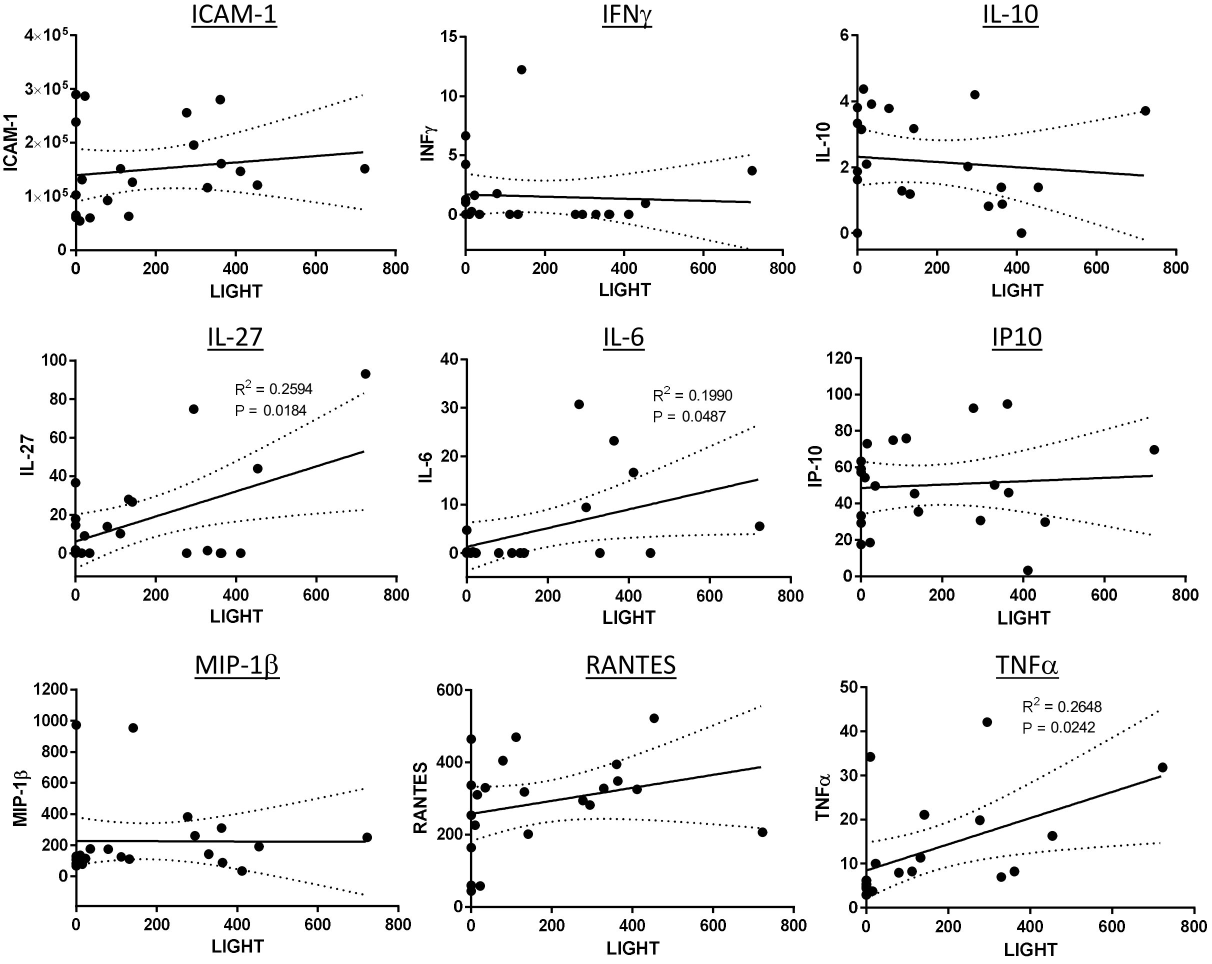


#### ‘Acute’ patients

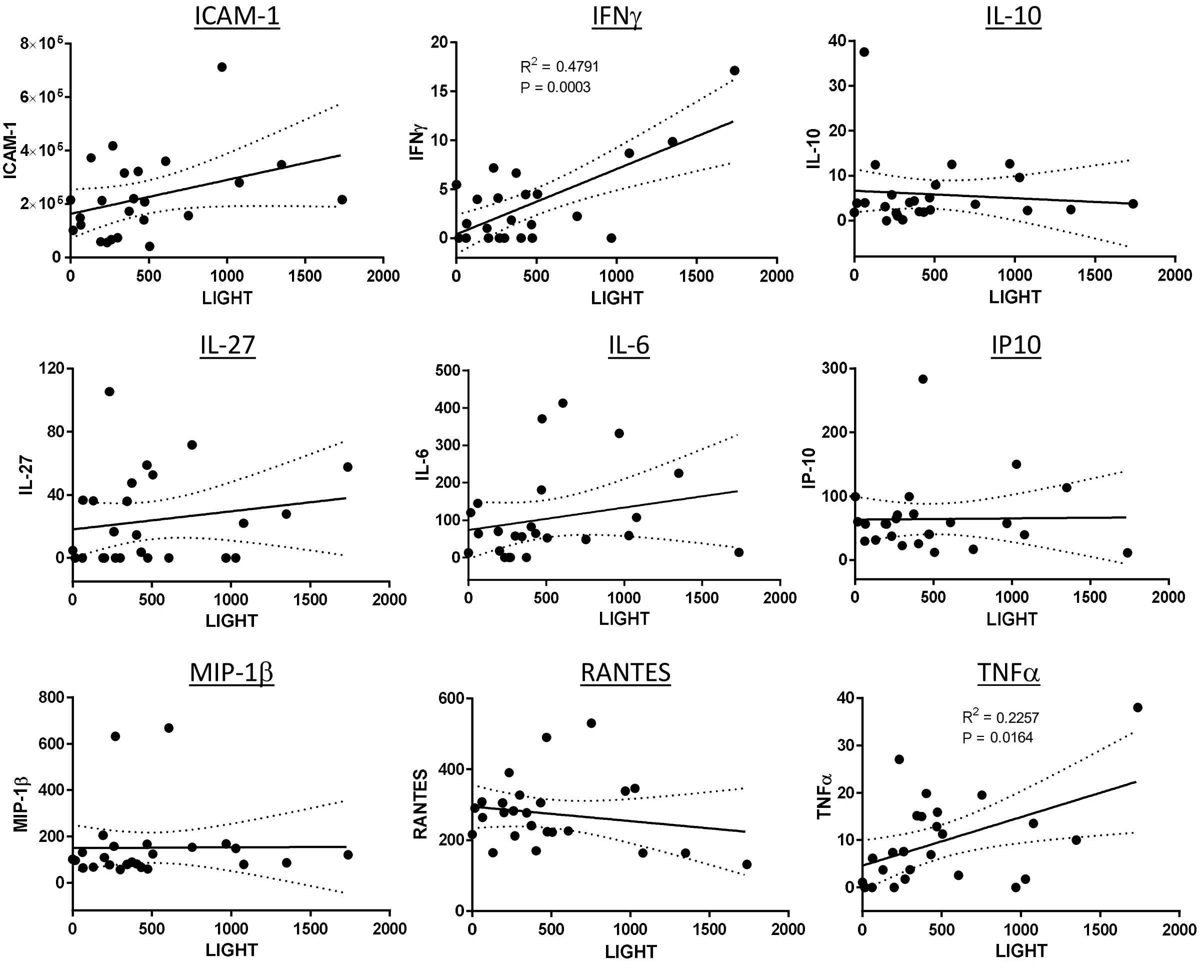


Supplemental figure S2

### Supplemental fig. S3

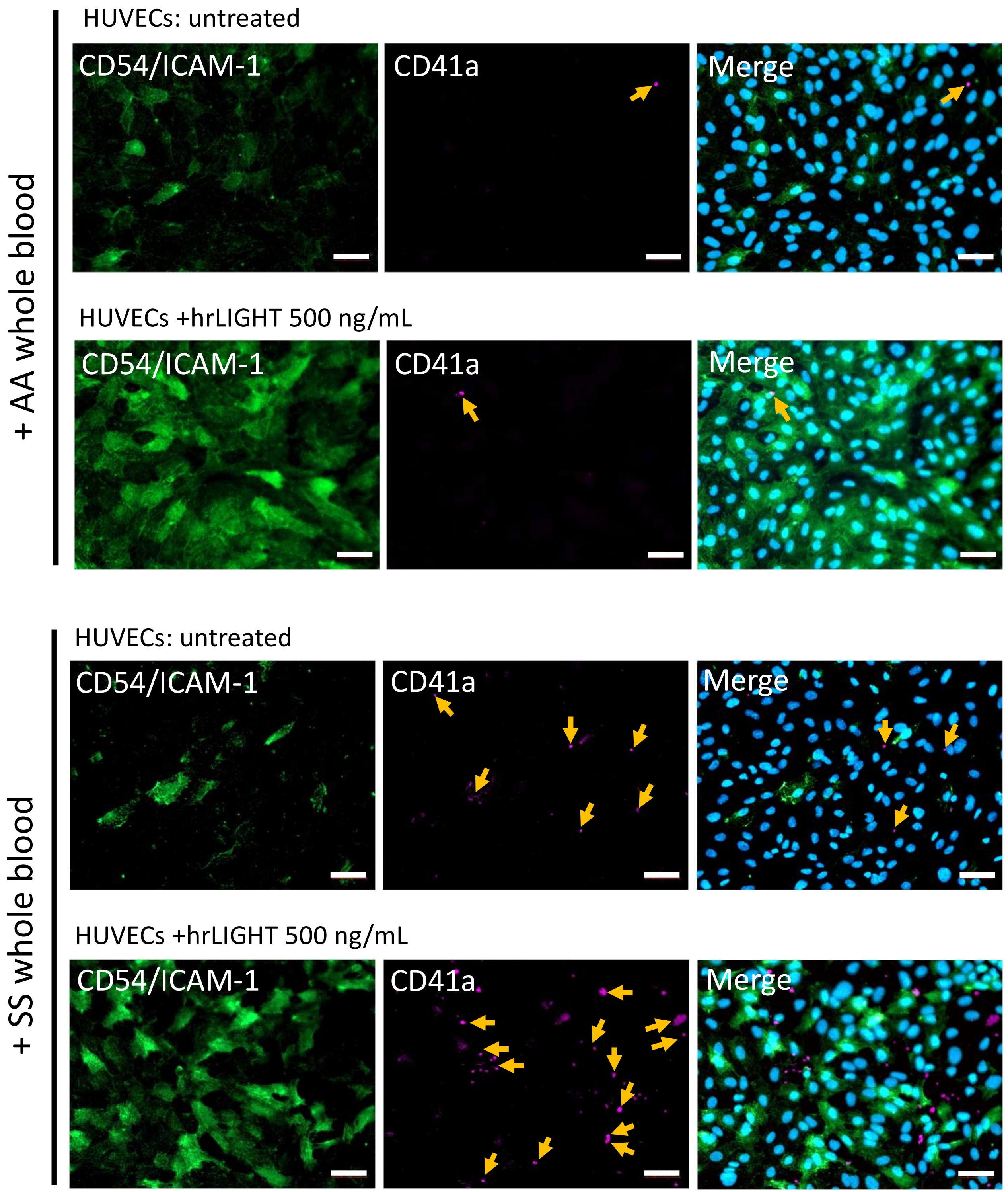


Supplemental fig. S3

### Supplemental fig. S5

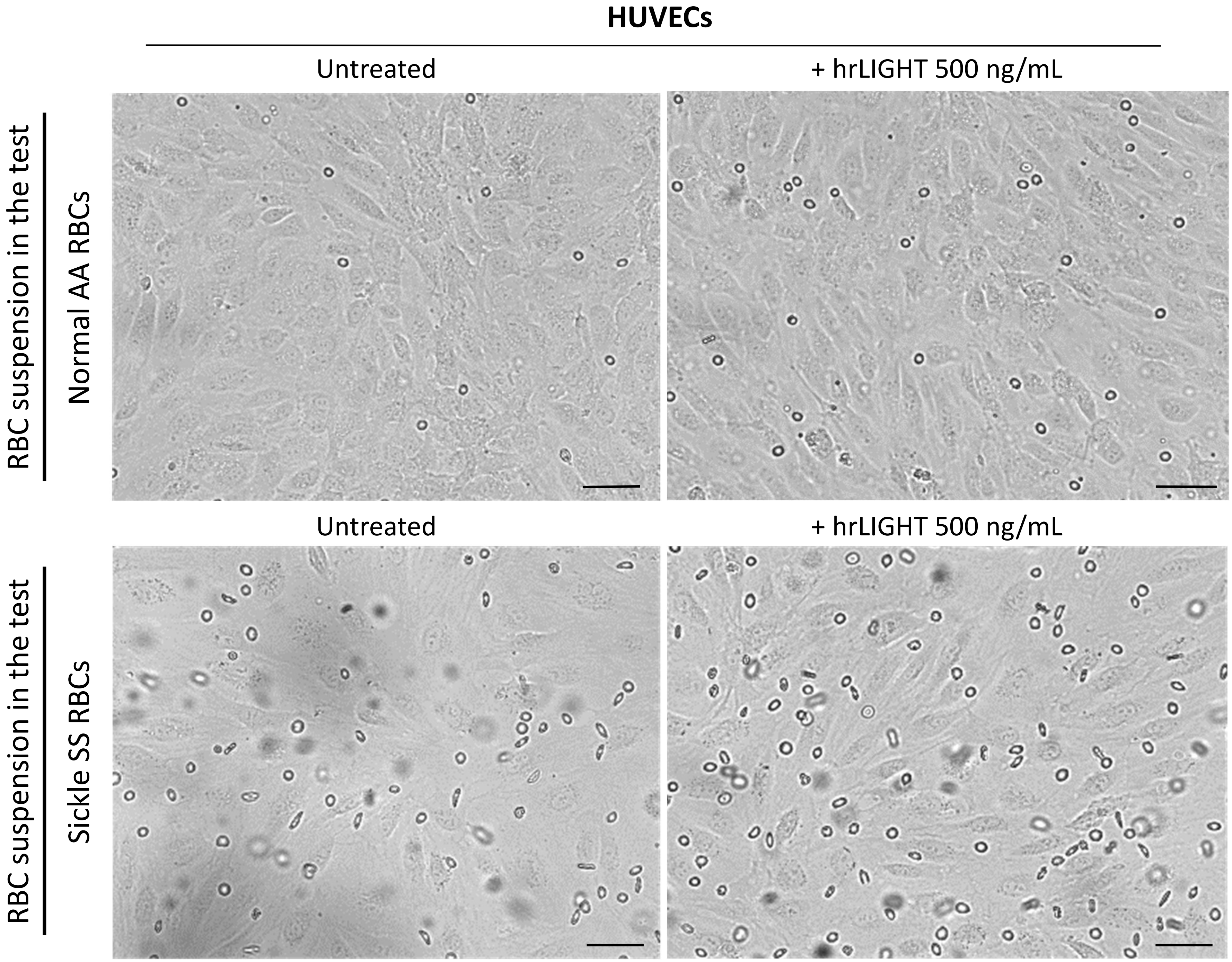


Supplemental figure S5

### Supplemental fig. S6

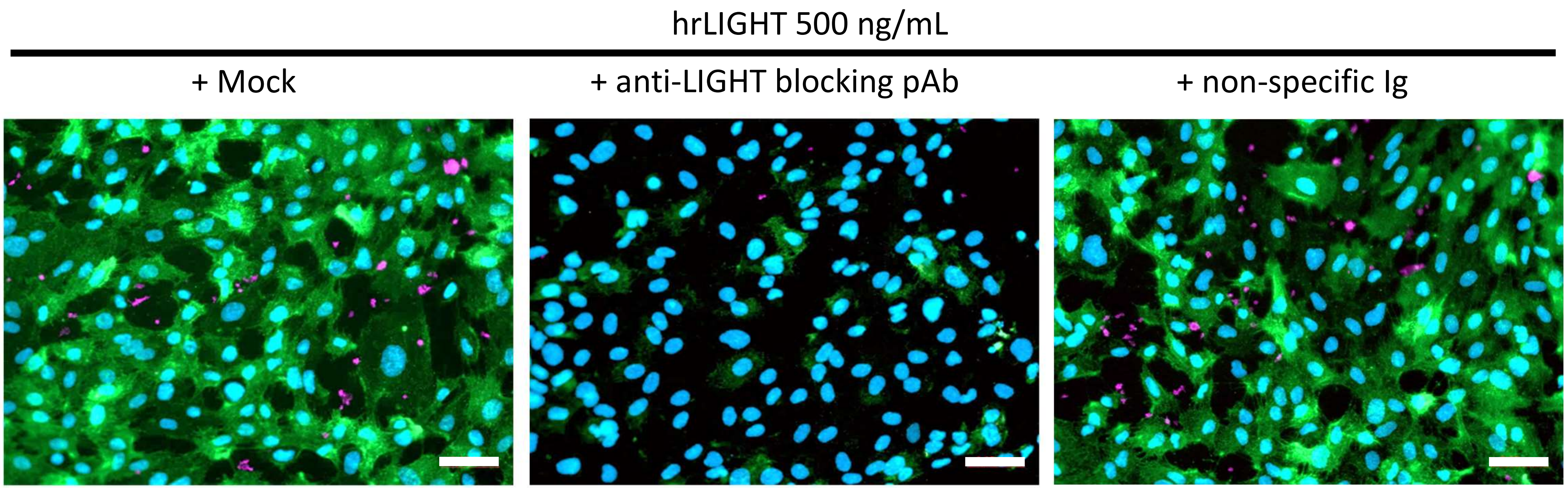


**CD54/ICAM-1** (green)/ **CD41a** (pink)/ **DAPI** (blue)

Magnification: 20x

Supplemental fig. S6
